# Confronting preferential sampling in wildlife surveys: diagnosis and model-based triage^†^

**DOI:** 10.1101/080879

**Authors:** Paul B. Conn, James T. Thorson, Devin S. Johnson

## Abstract

**Summary:**

1. Wildlife surveys are often used to estimate the density, abundance, or distribution of animal populations. Recently, model-based approaches to analyzing survey data have become popular because one can more readily accommodate departures from pre-planned survey routes and construct more detailed maps than one can with design-based procedures.
2. Species distribution models fitted to wildlife survey data often make the implicit assumption that locations chosen for sampling and animal abundance at those locations are conditionally independent given modeled covariates. However, this assumption is likely violated in many cases when survey effort is non-randomized, leading to preferential sampling.
3. We develop a hierarchical statistical modeling framework for detecting and alleviating the biasing effects of preferential sampling in species distribution models fitted to count data. The approach works by jointly modeling wildlife state variables and the locations selected for sampling, and specifying a dependent correlation structure between the two models.
4. Using simulation, we show that moderate levels of preferential sampling can lead to large (e.g. 40%) bias in estimates of animal density, and that our modeling approach can considerably reduce this bias.
5. We apply our approach to aerial survey counts of bearded seals (*Erignathus barbatus*) in the eastern Bering Sea. Models that included a preferential sampling effect led to lower estimates of abundance than models without, but the effect size of the preferential sampling parameter decreased in models that included explanatory environmental covariates.
6. When wildlife surveys are conducted without a well-defined sampling frame, ecologists should recognize the potentially biasing effects of preferential sampling. Joint models, such as those described in this paper, can be used to test and correct for such biases. Predictive covariates are also useful for bias reduction, but ultimately the best way to avoid preferential sampling bias is to incorporate design-based principles such as randomization and/or systematic sampling into survey design.

## Introduction

Surveys of unmarked animal populations are often used to estimate abundance and occurrence of animal populations and to predict species distributions, enterprises central to conservation, ecology, and management. For studies of abundance, researchers historically relied on design-based statistical inference (e.g. Cochran 1977), which requires adoption of a pre-defined sampling frame (e.g. using systematic random sampling, stratified random sampling, or some variant thereof). Designing animal surveys is relatively straightforward in such applications, and unbiased point and variance estimators are available. Recently, however, there has been a surge in research describing model-based procedures for estimating abundance, density, and occupancy from surveys of unmarked animals, including N-mixture and Dail-Madsen models for repeated point counts (Royle 2004; Dail & Madsen 2011), occupancy models for presence-absence surveys (MacKenzie *et al.* 2002; Johnson *et al.* 2013), and various model-based formulations for distance-sampling data (Hedley & Buckland 2004; Johnson *et al.* 2010; Miller *et al.* 2013). In such applications, it is common to use habitat or environmental covariates together with spatial effects (e.g. via trend surfaces or spatial random effects) to predict density or distributions across the landscape. We shall refer to the amalgam of model-based approaches for making spatially explicit inference about animal populations as “species distribution models” (SDMs; *sensu* Elith & Leathwick 2009), even though this term is more often used to refer to animal occurrence than it is to density or abundance.

One of the main advantages of using SDMs is that one is no longer beholden to predetermined sampling frames, and can potentially use data gathered from non-randomized designs or platforms of opportunity to make inferences about animal populations (Johnson *et al.* 2010). However, in a recent paper, Diggle *et al.* (2010) emphasized that spatially explicit statistical models can easily provide biased estimates when sampling disproportionately targets locations where the response of interest is higher (or lower) than expected given a particular set of explanatory covariates. In the context of SDMs, this might occur if sampling disproportionately occurs in locations where animals are known to be present or of high abundance. For example, if volunteer inventory participants have access to multiple sites with similar covariate values, bias might arise if they consistently choose sites where species are thought or known to be present. Bias might also arise if surveying effort is higher near bases of operations, and if animal abundance is higher (or lower) near bases of operations than elsewhere in the landscape.

In this article, we explore potential for bias in SDMs resulting from preferential sampling (hereafter, PS), and describe several model-based approaches for detecting and correcting for such biases. We start by describing a common currency for notation and basic model structures considered in this paper. Second, we review PS bias in a mathematical light, and describe prior approaches to coping with its effects. Third, we introduce a novel generalization of previously proposed PS models, allowing the investigator to jointly model animal encounter data and the locations chosen for sampling, including possible dependence structure between these two types of observations. Fourth, we conduct a simulation study to examine the performance of traditional SDMs and our newly developed PS model when data are gathered preferentially. Finally, we demonstrate our modeling approach by analyzing aerial survey counts of bearded seals (*Erignathus barbatus*) in the Bering Sea.

## Materials and methods

### NOTATION AND BASIC MODEL STRUCTURES

We focus here exclusively on discrete space (areal) models for animal encounter data as these seem to be the dominant form used in design and analysis of animal population surveys, although we note that PS is likely to affect analyses similarly regardless of the choice of spatial domain. We suppose that the investigator intending to fit a SDM to animal encounter data breaks their study area up into *S* survey units (label these *U*_1_, *U*_2_,…,*U*_*S*_), of which *n* are selected for sampling (call the set of sampled locations 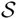). Each survey unit *i* is assigned a vector of covariates, **x**_*i*_, and an indicator *R*_*i*_ that takes on the value 1.0 if location *i* is sampled (i.e. if 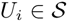), and is 0 otherwise. To formulate a “traditional” SDM, one could then write animal abundance or occurrence as a stochastic realization of a probability mass function *f*():

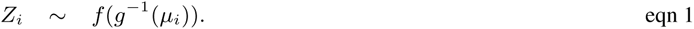

In this example, *Z*_*i*_ denotes the state variable of interest (e.g. occupancy or abundance), *g*() is a link function (e.g. probit or logit for occupancy, log for count data), and *µ*_*i*_ is a link-scale intensity value. In applications described in this paper, we write the intensity as

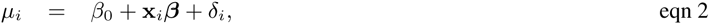

where *β*_0_ is an intercept parameter, **x**_*i*_ is a row vector of *m* predictive covariates associated with site *i*, ***β*** = {*β*_1_, *β*_2_,…, *β*_*m*_} is a column vector of *m* regression parameters, and ***δ*** = {*δ*_1_, *δ*_2_, …, *δ*_*S*_} are spatially autocorrelated random effects. For occupancy, *f*() would typically be Bernoulli, while the Poisson or negative binomial are typically choices for analysis of count data; common forms for *δ*_*i*_ include geostatistical specifications (Cressie 1993; Diggle *et al.* 1998), Gaussian Markov random fields (e.g. conditionally autoregressive models; Rue & Held 2005), or low rank alternatives such as predictive process (Banerjee *et al.* 2008; Latimer *et al.* 2009) or restricted spatial regression models (Reich *et al.* 2006; Hughes & Haran 2013).

The model for *Z*_*i*_ describes variation in the process of interest and is often described as the “process” model. However, it is usually impossible to observe the system perfectly even in locations where sampling occurs, so it is customary to include an observation model describing incomplete detection. For occupancy studies, the response variable *Y*_*i*_ = 1 if the species of interest is detected and is 0 otherwise, and is modeled with a Bernoulli distribution (Royle & Dorazio 2008):

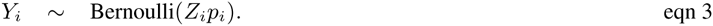

Here, the detection probability *p*_*i*_ is possibly a function of survey and observer specific covariates. Replicate surveys of the same sampling unit provide the necessary information to estimate *p*_*i*_. For count surveys, a possible model is

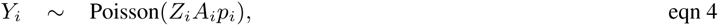

where the *Y*_*i*_ now represents the count of animals obtained while surveying unit *i*, *A*_*i*_ denotes the proportion of sample unit *i* that is surveyed, and *p*_*i*_ gives detection probability. Additional information will often be needed to estimate *p*_*i*_ in this context, such as data from double observers, distance observations, or double sampling (see e.g. Buckland *et al.* 2001; Royle *et al.* 2004; Borchers *et al.* 2006; Conn *et al.* 2014).

For the remainder of this treatment, we use bold symbols to denote vector-valued quantities or matrices. We also use standard bracket notation to denote probability mass and density functions. For instance [**Z**] denotes the marginal probability mass function for **Z**, and [**Z**|**Y**] represents the conditional distribution of **Z** given **Y**. We use ***µ*** and ***ν*** to denote log-scale abundance and the logit of the probability of sampling, so that *Z*_*i*_∼ *f*(*µ*_*i*_), and *R*_*i*_∼ *f* (*ν*_*i*_). We use the notation *Z*_*i*_ when describing the state process in general terms, but often switch to the conventional notation *N*_*i*_ when animal abundance is the explicit focus of interest.

### PREFERENTIAL SAMPLING: A PRIMER

One of the appealing aspects of model-based estimation is that there is no requirement that surveys rely on a pre-planned survey design selected probabilistically from an underlying sampling frame. For instance, investigators can reallocate sampling effort if weather or logistics preclude surveying in a desired location. This can be a crucial advantage in surveys covering large areas with frequent inclement weather. It also opens the door for using platforms of opportunity, presence only, and citizen science data for estimation.

However, the manner in which effort is ultimately allocated can potentially have profound influence on SDM estimator performance. With respect to nonrandom sampling, two possible problems seem particularly likely in discrete spatial domains: coarse scale preferential sampling (CSPS), and fine scale preferential sampling (FSPS) (Fig. 1). FSPS arises when the observations taken at a particular sampling unit are non-random with respect to the density of animals within that sampling unit. For instance, when allocating line transect survey effort, it may be tempting to place the transect in a manner that targets habitat or landscape features that maximize the number of animals that will be encountered. However, this strategy will clearly lead to positive bias when estimating density or abundance.

**Fig. 1.**
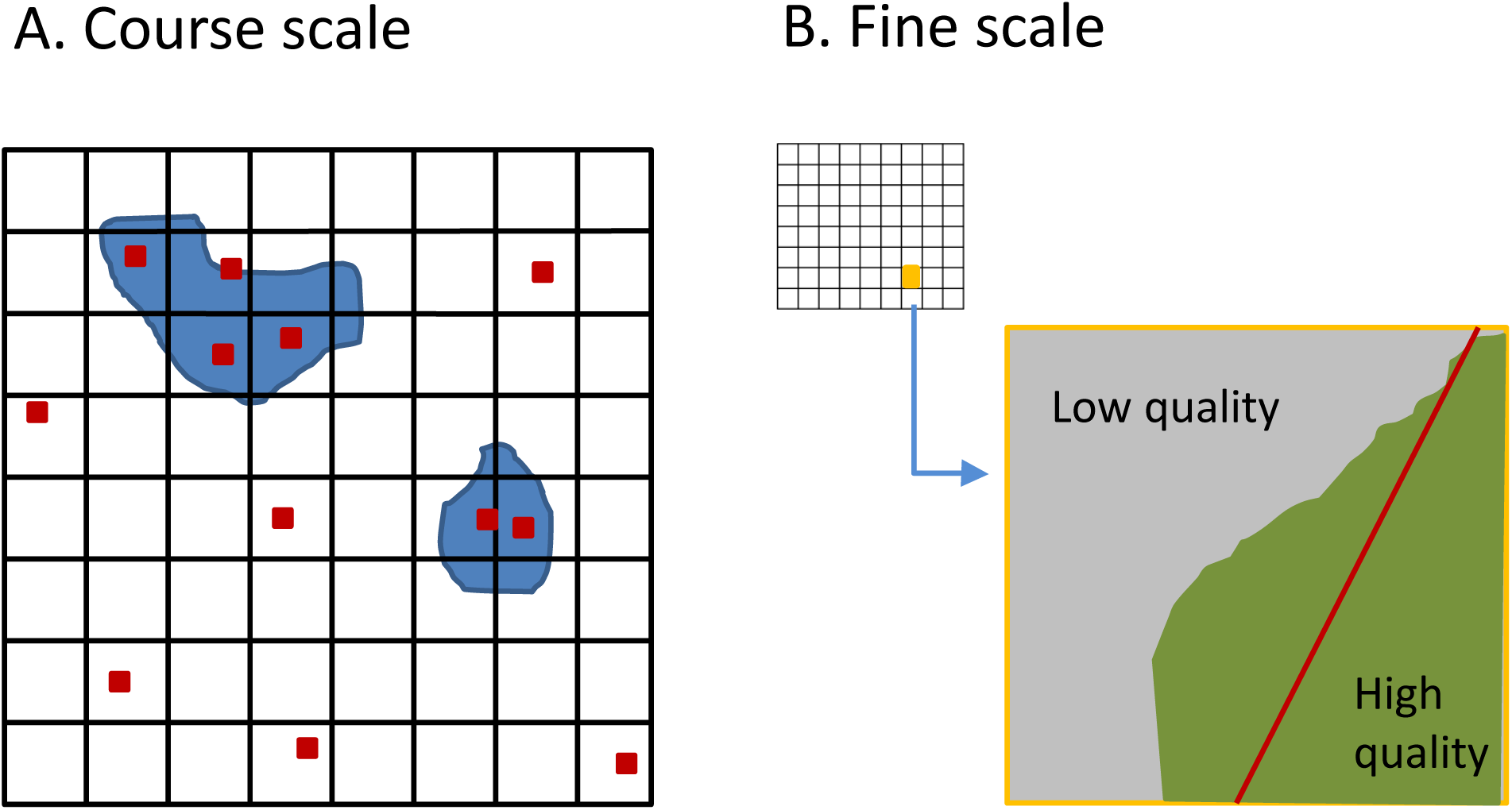
A depiction of two types of preferential sampling. In (A), an investigator preferentially places point transects (red squares) within regions of high known animal density (blue polygons). This can cause bias in abundance or occupancy estimators unless this a priori knowledge about density is explicitly modeled. In (B), a fine scale version of preferential sampling occurs when a line transect (red line) is intentionally placed across a region of high quality habitat. If a landscape is discretized into homogeneous survey units for analysis (as in a grid), it is essential that the habitat surveyed within each survey unit be randomly determined when estimating abundance. If not, bias (usually positive) can be expected.

By contrast, CSPS (hereafter, PS), the primary focus of this article, arises when the locations being sampled and the process of interest (e.g. density, occupancy) are conditionally dependent given modeled covariates (Diggle *et al.* 2010). For instance, PS can occur when the investigator uses a priori knowledge or observations of the state variable obtained during sampling to allocate survey effort in places where abundance or occurrence is known to be high. Diggle *et al.* (2010) showed that this type of PS can lead to bias when this extra information is not included in models for the state variable of interest. Specifically, PS arises when we consider the set of sampled locations as stochastic and when [**R**, **Z**|**x**] ≠ [**R**|**x**][**Z**|**x**], where **R** is an indicator vector whose elements *R*_*i*_ are 1.0 if sampling unit *i* is sampled and are zero otherwise. We use this definition of PS throughout the rest of the manuscript, noting that it is somewhat different than has sometimes been used in the SDM literature. For instance, Merckx *et al.* (2011) use the term “preferential sampling” to refer to the process of visiting some sites more often than others, while Manceur & Kühn (2014) define it as occurring when the locations selected for sampling are a function of an environmental covariate. Neither of these latter conditions are problematic outside of the specialized field of presence-only modelling.

Diggle *et al.* (2010) demonstrated PS with an environmental monitoring problem, whereby pollutant monitoring stations were more highly clustered around urban areas with high concentrations of pollutants than in rural areas with comparably low levels of pollutants. Fitting simple geostatistical models without fixed effects led to positively biased estimates of landscape-level pollutant concentrations. Presumably (and as noted by discussants of the article) including a fixed effect associated with a relevant covariate (e.g. a development index) would likely reduce or eliminate bias. However, the primary point of Diggle *et al.* (2010) is well taken: inclusion of spatially autocorrelated random effects in a statistical model is insufficient to remove the potentially biasing effects of PS.

As in the pollution example, having good explanatory covariates may also reduce bias when fitting SDMs to animal encounter data under PS. However, in many ecological applications, predictive covariates explain only a small portion of variation present in the data. If the locations selected for sampling are a function of some unmodelled factor related to abundance (intentionally or unintentionally), bias may still occur. Despite the clear potential for bias in SDMs, there are few examples where PS (*sensu* Diggle *et al.* 2010) is discussed with regard to SDMs. One exception is Chakraborty *et al.* (2010), who acknowledged the likely presence of PS when fitting SDMs to data obtained using nonrandomized designs. However, they did not attempt to account for PS in their models.

In design-based sampling, unequal sampling intensity is often accommodated via stratification or unequal probability sampling, as with Horvitz-Thompson-like estimators where the probability of inclusion varies by sampling unit (Cochran 1977). However, in the case of PS, this inclusion probability also depends on the value of the response associated with the sampling unit. Evidently, any approach to account for PS should also account for the dependence between the state variable of interest and the locations chosen for sampling.

Several authors have attempted model-based corrections for PS in the statistical literature. For Gaussian models in a continuous spatial domain, Diggle *et al.* (2010) and *Pati et al.* (2011) jointly modeled the locations that are chosen for sampling and the underlying random field of interest. In particular, they expressed sampled locations as an inhomogeneous Poisson point process where the underlying log-scale intensity depended linearly on spatially-referenced random field values. For instance, writing observations of the spatial random field at a location *i* as

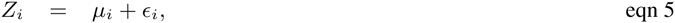

the spatially continuous relative intensity (*ψ*_*i*_) of sampling locations at *i* could be written as

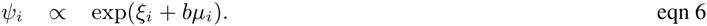

Here, the parameter *b* describes the level of PS; *b* = 0 implies no PS, *b* > 0 implies a greater level of sampling in locations where the spatial process (e.g. animal density) is high, and *b* < 0 implies greater sampling where the spatial process is low. Importantly, when explanatory covariates are used in models for *µ*_*i*_ and *ξ*_*i*_, Pati *et al.* (2011) show that “… accounting for informative sampling is only necessary when there is an association between the spatial surface of interest and the sampling density that cannot be explained by the shared spatial covariates.” Pati *et al.* (2011) also consider a simpler, plug-in based estimator, where the log of a nonparameteric estimate of sampling density (specifically, a two dimensional kernel density estimate) is used as an additional fixed effect in Eq. 5, finding that this approach helped reduce bias associated with PS, but did not perform as well as the full joint model.

### A GENERALIZED PREFERENTIAL SAMPLING MODEL

The models considered by Diggle *et al.* (2010) and Pati *et al.* (2011) are a useful first step in addressing and modeling PS. However, they are somewhat limited since they are specific to continuous spatial domains, continuous data (as opposed to presence/absence or count data), and Gaussian error distributions. Also, they require the linear predictor of the PS model to be written as a simple linear function of the the spatial process model for density. In real world applications, we can envision cases where sampling is strongly preferential in certain areas of the landscape, and not in others. For instance, sampling may be more strongly preferential close to bases of operations, (e.g. landing strips in the case of aerial surveys), but less so in areas that are harder to get to.

Given these limitations, our present task is to generalize PS models to the types of data more typical of SDMs, and to allow the degree of PS to vary across the landscape. Like Diggle *et al.* (2010) and Pati *et al.* (2011), we impose a joint model for the process of interest (animal abundance or occurrence) and the locations chosen for sampling. For the abundance process model, we start with eq. 1 as a general formulation for non-Gaussian data, writing the link-scale expectation as in eq. 2. Next, recalling that *R_i_* is a binary indicator taking on the value 1.0 if survey unit *i* is selected for sampling, and is 0.0 otherwise, we model *R*_*i*_ using a Bernoulli distribution:

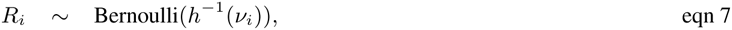

where *h*() denotes a link function appropriate for binary data (e.g. logit, probit). We then write the intensity for this model as

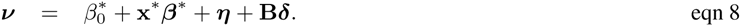

In a similar fashion to the model for the state process, the sampling intensity model has an intercept 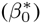, explanatory covariates 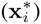, fixed effect regression parameters (***β***^∗^) and spatially autocorrelated random effects (***η*** and ***δ***). The predictive covariates **x**_*i*_ from Eq. 2 and 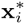 from Eq. 8 may or may not be the same. Note also that the spatially autocorrelated random effects ***δ*** are included in both Eqs. 2 and 8, allowing for dependency in the two models, with the matrix **B** describing the strength and type of dependence between the sampling process and underlying density. The spatially autocorrelated random effects ***η*** are assumed independent of the ***δ***. In practice, we find we often need to fix 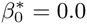 when random effects in Eq. 8 are estimated to permit parameter identification. The formulation in Eq. 8 is similar to the one previously proposed for hierarchical multivariate models with spatial dependence (cf. Royle & Berliner 1999). There are multiple ways of structuring **B** depending on the complexity of spatial dependence desired for the PS process (Royle & Berliner 1999). For instance, setting **B** = **0**_*S×S*_ corresponds to an absence of spatial dependence (and thus no PS). Setting **B** = *b***I**, where *b* is an estimated parameter and **I** is an (*S × S*) identity matrix corresponds to the linear PS model suggested by Diggle *et al.* (2010) and Pati *et al.* (2011). Alternatively, we could allow the degree of PS to vary across the landscape. For instance, one can contemplate a trend surface model for PS by specifying a diagonal matrix for **B**, with entries given by *b*_0_+ *b*_1_lat_*i*_+ *b*_2_long_*i*_, where *b*_0_, *b*_1_, and *b*_2_ are estimated parameters and lat_*i*_ and long_*i*_ give latitude and longitude, respectively (Royle & Berliner 1999). Theoretically, one could include more highly parameterized structures for spatial dependence, such as higher order trend surface or spline formulation (Royle & Berliner 1999), but the ability to robustly estimate the parameters of such a model is likely dependent on having a rich, spatially balanced dataset, which is often not the case in ecological applications.

A comparison of the performance of models with different sets of constraints on **B** can serve as a test of PS. In particular, if one can demonstrate that models with **B** = **0** perform similarly or better than models with **B** ≠ **0**, then PS is likely not worth modeling and inference can proceed using standard SDMs (i.e. not modeling sampling intensity).

### SIMULATION STUDY

To illustrate PS and demonstrate that our proposed model has reasonable performance, we conducted a small simulation experiment. For each of 500 simulations, we generated abundance of a hypothetical species over a 25 *×* 25 grid as

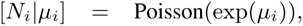

where *i* indexes survey unit *i*, and *µ*_*i*_ is determined according to Eq. 2. Abundance was generated as a function of a single spatially autocorrelated landscape covariate, as well as residual spatial autocorrelation (*δ*_*i*_) and overdispersion (fig. 2). Specific details of data generation procedures are provided in Appendix S1.

**Fig. 2.**
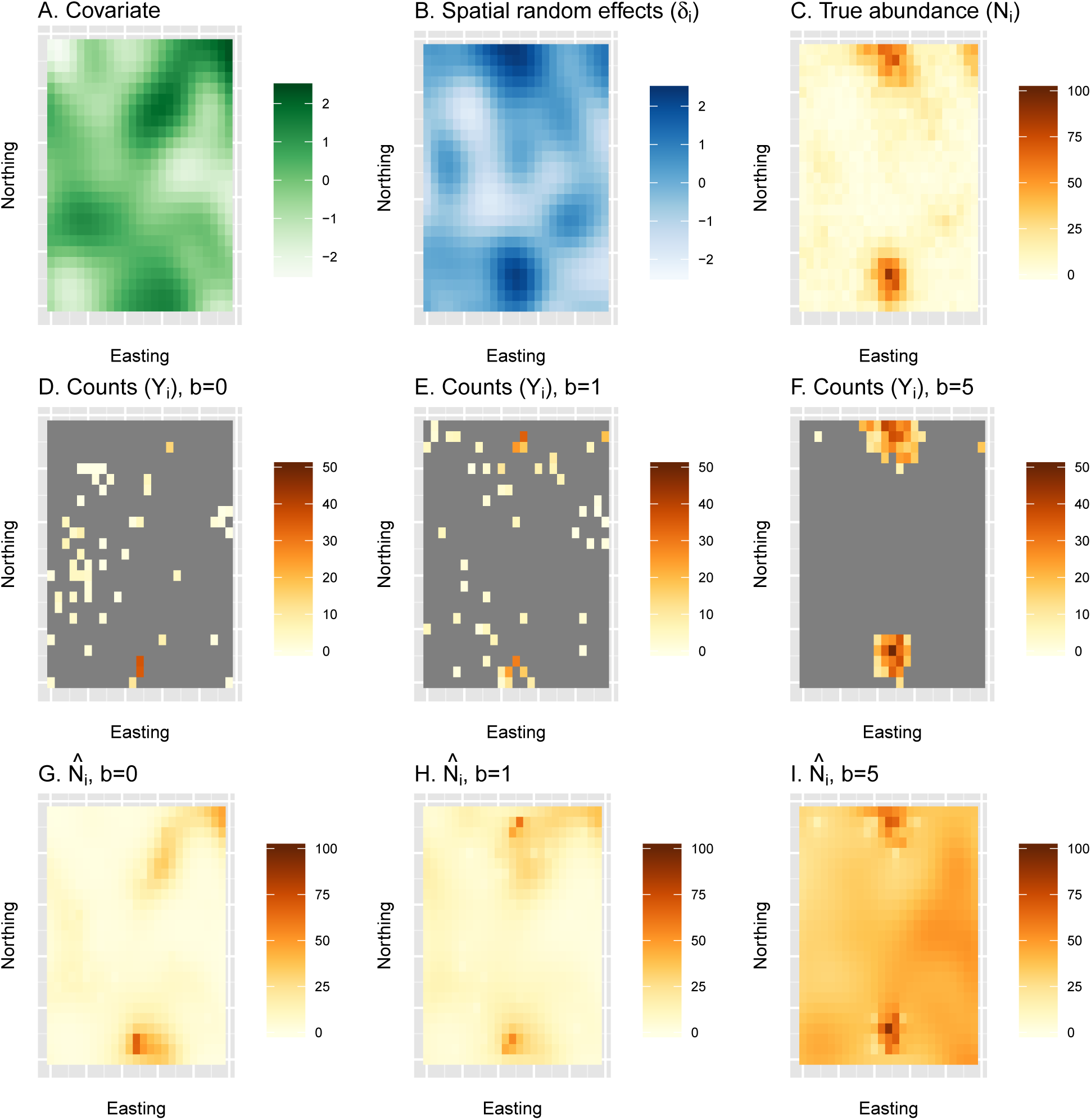
An example of a single simulation replicate examining estimates of abundance from a naive species distribution model under preferential sampling. First, true abundance (C) is generated as a function of a spatially autocorrelated covariate (A) and a spatially autocorrelated random effect (B). Second, counts are generated for three different types of surveys, including a simple random sample (*b* = 0; D) and surveys with moderate (*b* = 1; E) or pathological (*b* = 5; F) levels of preferential sampling. Finally, spatially explicit estimates of abundance are generated using a traditional SDM (with *b* set to 0.0) to each of the count datasets (G-I). In this particular simulation replicate, cumulative abundance was underestimated by 18% when *b* = 0, overestimated by 17% when *b* = 1, and overestimated by 293% when *b* = 5. For a summary of bias over 500 simulation replicates, see fig. 4.

For each simulated landscape we generated three virtual count surveys using eqs. 7 and 8. Each survey had ***β**^∗^* = *η*_*i*_ = 0 (that is, no covariate or spatially autocorrelated random effects), but differed in how the matrix **B** was parameterized. In the first, we set **B** = 0, so that surveyed locations were selected independently of the abundance generating process. For the second and third, we set **B** to be a diagonal matrix with entries *b* = 1 and *b* = 5, respectively, so that the probability of sampling a given survey unit (grid cell) was explicitly dependent on the latent abundance in that unit. We refer to these scenarios as moderate and pathological PS, respectively (see fig. 3). Simulations were configured so that *n* = 50 of the 625 survey units were sampled; each survey was set to cover half of the target cell.

**Fig. 3.**
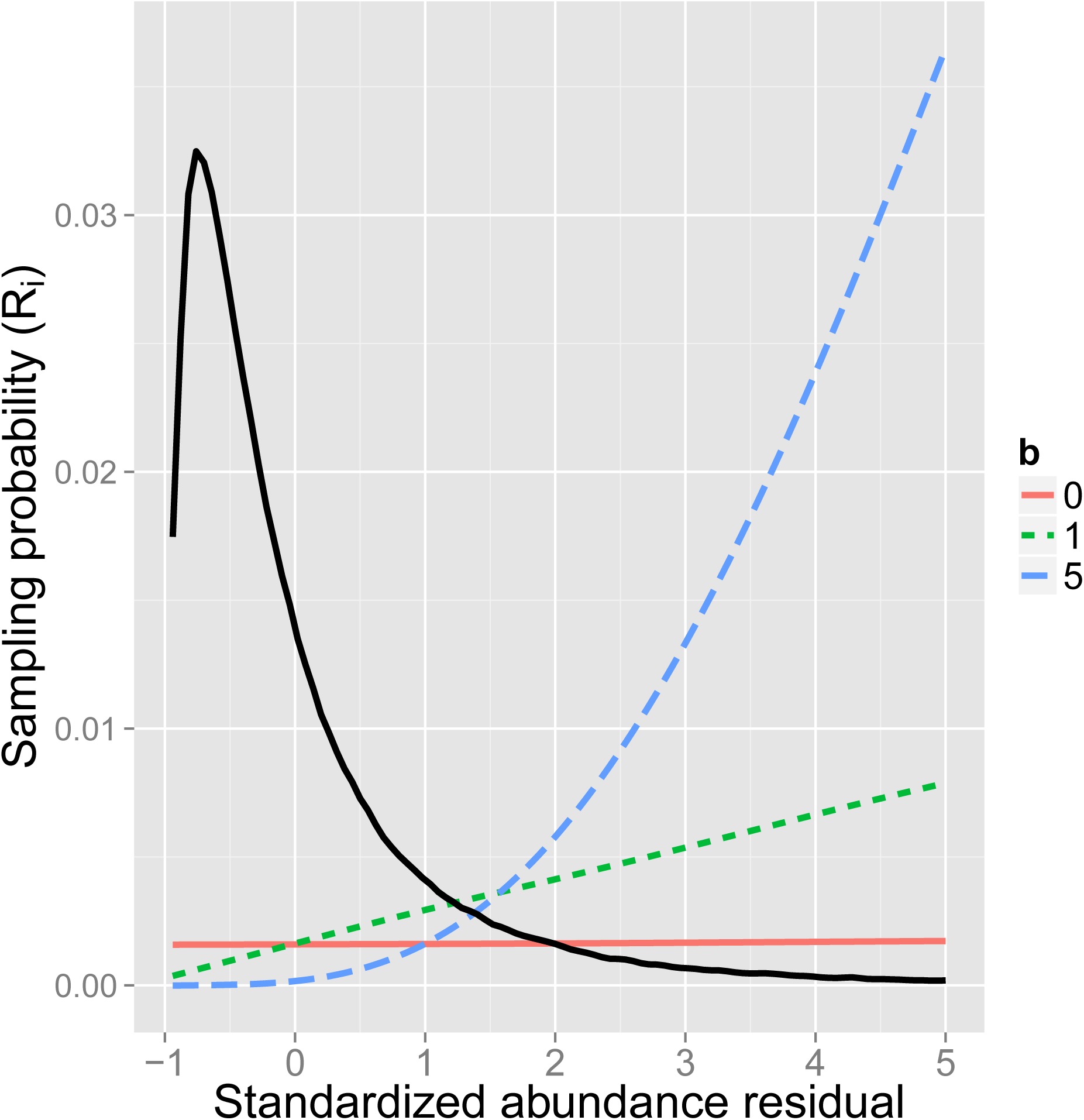
Expected relationship between the probability of a survey unit being selected for sampling and its abundance residual in the simulation study. The base case *b* = 0 represents simple random sampling, while *b* = 1 and *b* = 5 represent moderate and pathological levels of preferential sampling, respectively. Also shown are is the realized distribution (smoothed histogram) of abundance residuals among survey units in the simulation study, scaled to fit in the plot margins (solid black line).

We fitted two different models to each count dataset, both of which were provided the habitat covariate used (in part) to generate the data for which a log-linear coefficient *β* was estimated. In the first model, the elements of **B** in eq. 8 were all set to zero. In this case, the abundance and sampling process submodels were independent, as is the case canonical SDMs (at lest when fitted to presence-absence or count data). In the second model, we included an explicit connection between the distribution of animal abundance and the sampling process by setting **B** = *b***I**, where *b* is an estimated parameter, and **I** is an identity matrix.

We used maximum likelihood to conduct statistical inference. In particular, we used Template Model Builder (TMB; Kristensen *et al.* 2016), interfaced with the R programming environment, to conduct maximization. The TMB software uses a Laplace approximation to integrate out random effects (***η*** and ***δ***), and a bias correction algorithm (Tierney *et al.* 1989; Thorson & Kristensen In Press) to obtain abundance estimates and standard errors that properly account for nonlinear transformations of random effects. This approach resulted in a facile implementation and speedy computing times, allowing us to conduct simulation and model testing with greater efficiency than would have been possible with Bayesian simulation. Further detail on statistical methods are provided in Appendix S1; requisite R and TMB code will be published to a publicly accessible repository upon acceptance, and is also available at https://github.com/NMML/pref_sampling/.

### BEARDED SEAL COUNT SURVEYS

We applied our modeling technique to counts of bearded seals obtained on aerial transects flown over the eastern Bering Sea from 10-16 April, 2012 (Fig. 5). These counts were gathered as part of a larger survey designed to estimate abundance of four species of ice-associated seals; the survey is described in greater detail elsewhere (Conn *et al.* 2014, 2015). The survey area considered here consists of 25 by 25 km grid cells bordered to the north by the Bering strait, to the west by the international date line, to the south by maximal April ice extent, and to the east by the Alaska, USA mainland. Here, we limit counts to those gathered within a one week period so that relative abundance will remain relatively constant throughout the study area. Our primary focus in this application is to diagnose PS (rather than to estimate absolute abundance). As such, we do not attempt to correct for nuisance processes such as incomplete detection or species misclassification, which requires models of increased sophistication (Conn *et al.* 2014).

Our choice to model bearded seal counts, as opposed to one of the other seal species, is based on the observation that bearded seal densities tend to be highest in the northern portion of the study area. This is also the location of one of the primary airports used to prosecute surveys (Nome, Alaska, USA). Higher survey coverage in areas of high bearded seal density could potentially lead to positive bias in apparent abundance owing to PS.

To test for such an effect, we modeled bearded seal counts using the formulation

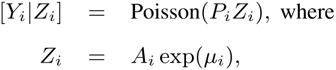

where *P*_*i*_ defines the proportion of grid cell *i* that is sampled, *A*_*i*_ gives the proportion of grid cell *i* that is composed of salt water habitat, and *µ*_*i*_ is defined in Eq. 2. We modeled the grid cells that were chosen for sampling using Eqs. 7–8.

We fitted a total of six models (*M*_*cov*=0,*b*=0_, *M*_*cov*=0,*b*=1_, *M*_*cov*=1,*b*=0_, *M*_*cov*=1,*b*=1_) to bearded seal count data using the same estimation framework as in the simulation study. Models varied by (i) whether or not habitat and landscape variables were used as predictors of bearded seal density (*cov* = 1 and *cov* = 0, respectively), and (ii) the form of PS (*b* = 0 indicates no PS; *b* = 1 indicates **B** = *b***I**, where *b* is an estimated parameter). We also attempted to fit models where the PS **B** matrix varied over the landscape using a trend surface specification, but parameter identification was suspect in these models and are not reported here (see Appendix S2 for more information). When habitat and landscape variables were included, we used three log-linear predictors: linear and quadratic functions of sea ice concentration, and distance from the southern ice edge. Remotely sensed sea ice data were obtained at a 25 *×* 25 km resolution from the National Snow and Ice Data Center, Boulder, CO, USA, as described by Conn *et al.* (2014). Models for *µ*_*i*_ and *ν*_*i*_ both utilized spatially autocorrelated random effects with a Matérn covariance function between grid cell centroids (see Appendix S1 for further details). When covariates were included, they were included in both models (i.e. for *µ*_*i*_ and *ν*_*i*_).

## Results

### SIMULATION STUDY

Estimates of cumulative animal abundance across simulated landscapes were median unbiased for both estimation methods when the sites selected for sampling were independent of animal density, though when *b* was estimated, abundance estimates were more right skewed and had higher variance (fig. 4). Under moderate PS (*b* = 1), estimation of the PS parameter *b* led to a median bias of 5%, while the canonical SDM model ignoring preferential sampling had a median bias of 40%. Under pathological PS (*b* = 5), both estimation methods were extremely biased, but was even more severe for the naive model ignoring PS (fig. 4).

**Fig. 4.**
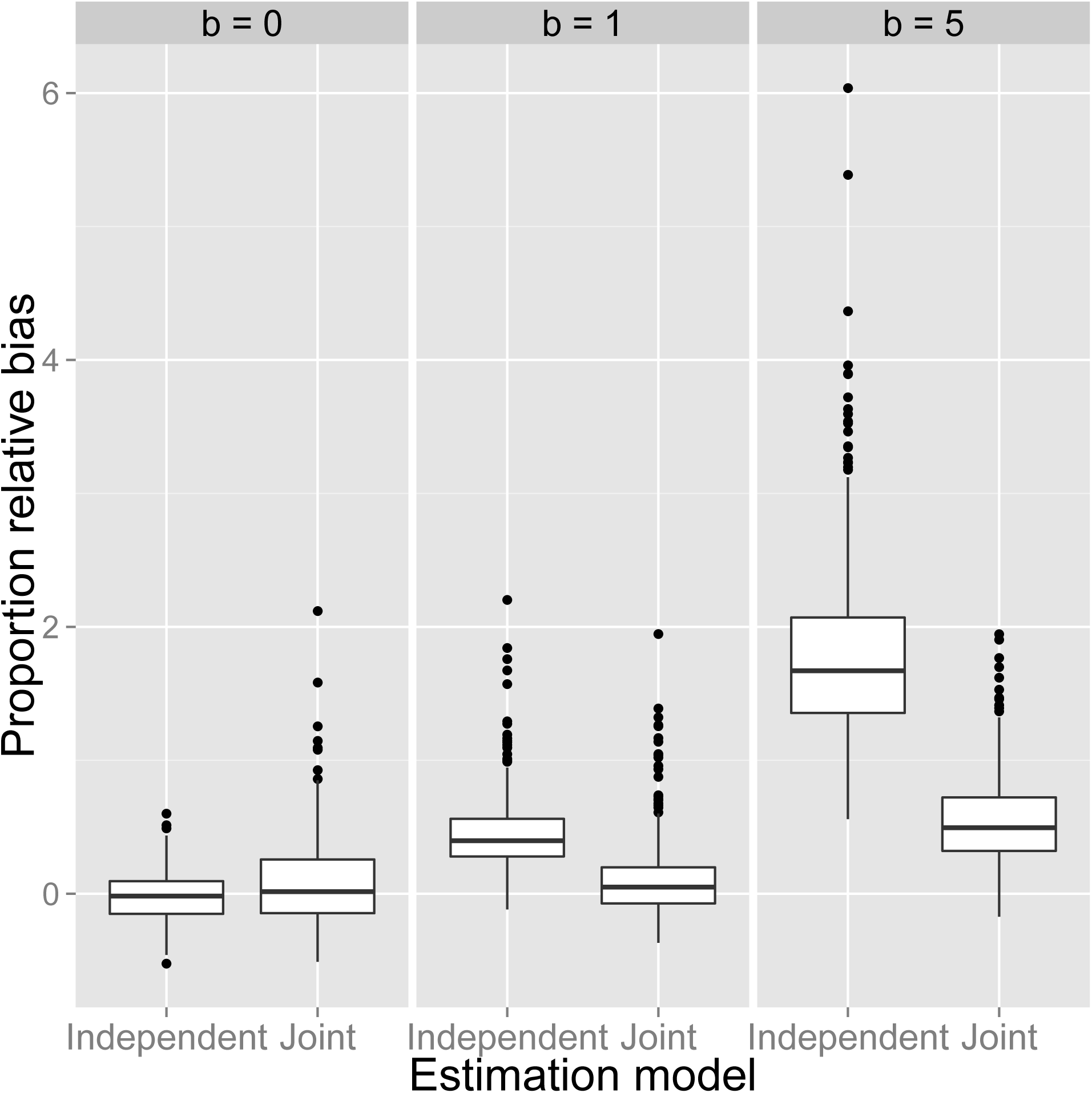
Relative proportional error in abundance from the simulation experiment as computed with respect the posterior mode with a bias correction. Each boxplot summarizes the distribution of relative proportional error as a function of the type of sampling, including simple random sampling (*b* = 0), moderate preferential sampling (*b* = 1), and pathological preferential sampling (*b* = 5). Results vary by the type of estimation model; in the “independent” model, *b* is set to 0.0; in the “joint” model, *b* is estimated. Lower and upper limits of each box correspond to first and third quartiles, while whiskers extend to the lowest and highest observed bias within 1.5 interquartile range units from the box. Points denote outliers outside of this range. Horizontal lines within boxes denote median bias.

**Fig. 5.**
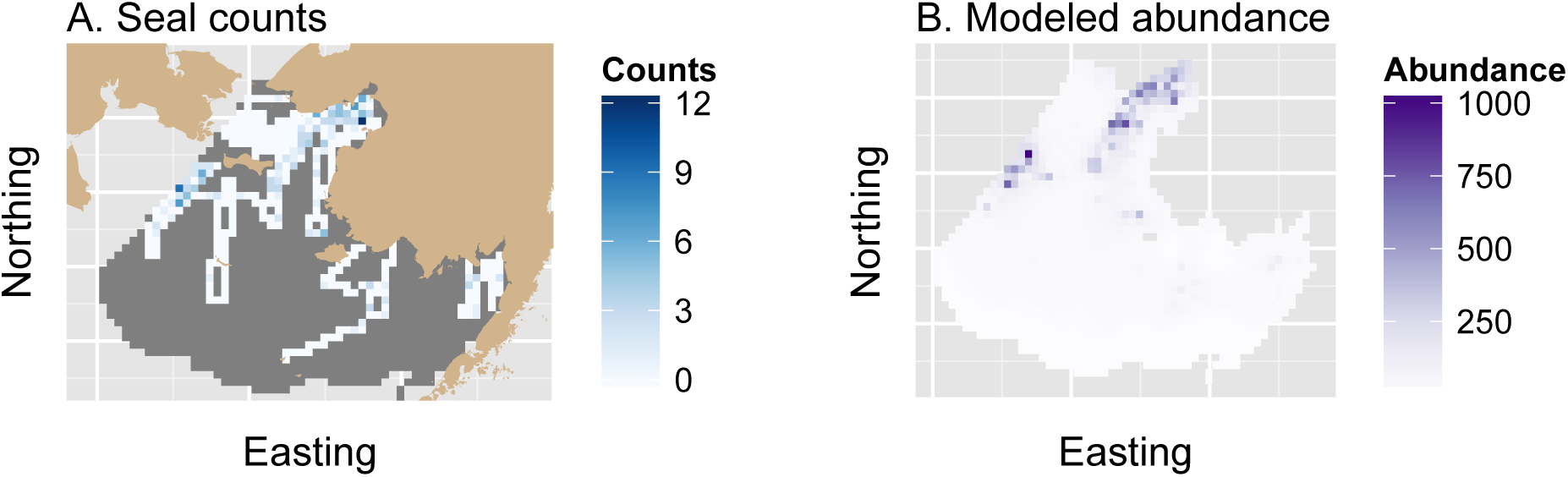
Aerial survey counts and estimated apparent abundance of bearded seals in the eastern Bering Sea, April 10-16, 2012. Counts and estimates are shown relative to a survey grid that extends south from the Bering Strait and borders the Alaska, USA mainland to the east. In (A), tan shading denotes land, unsurveyed grid cells appear in dark gray, and counts appear in a white-blue spectrum. Apparent bearded seal abundance estimates (B) are presented from the model with the lowest integrated AIC score, which included covariate effects but no preferential sampling effect. Apparent abundance estimates are uncorrected for imperfect detection or species misclassification.

### BEARDED SEAL ANALYSIS

Marginal AIC strongly favored models with covariate effects, but for such models the presence of PS was equivocal (Table 1). Further intuition can be gained by examining estimates of the PS parameter, *b*. For the PS model without predictive covariates, 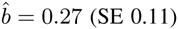, and for the model with predictive covariates, 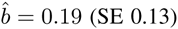. Thus, it appeared that including predictive covariates decreased the PS effect size, as suggested by Pati *et al.* (2011). Estimates of abundance were substantially higher for models without a PS effect, with the non-PS model having a 49% higher estimate when covariates were not modeled, and a 19% higher estimate when covariate effects were included.

**Table 1.** A summary of model selection results and estimated abundance for the four models fitted to bearded seal counts. The models include formulations with or without predictive covariates (*cov* = 1 or 0, respectively), and with or without the preferential sampling parameter *b* estimated (*b* = 1 or 0, respectively). All models included spatially autocorrelated random effects on log-scale abundance intensity. Shown are the log integrated likelihood, the number of fixed effect parameters, ∆AIC, AIC model weights, and estimated apparent abundance over the landscape 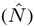 together with a Hessian-based standard error estimate.

| Model | Log likelihood | Params | $\Delta AIC$ | Wgt | $\hat{N}(\text{SE})$ |
| --- | --- | --- | --- | --- | --- |
| $M_{cov=0,b=0}$ | -2667.1 | 3 | 21.7 | 0.00 | 68556 (7408) |
| $M_{cov=0,b=1}$ | -2665.3 | 4 | 20.1 | 0.00 | 45857 (5114) |
| $M_{cov=1,b=0}$ | -2650.3 | 9 | 0.0 | 0.53 | 59312<sup>†</sup> (5231) |
| $M_{cov=1,b=1}$ | -2649.4 | 10 | 0.3 | 0.47 | 49826 (10369) |

<sup>†</sup> Refitted model; see *Results*.

Note that unlike the other models, *M*_*cov*=1,*b*=0_ predicted anomalously high bearded seal abundance in the extreme southern portion of the study area where sea ice was absent (where there was no habitat for seals). Thus, while we present original likelihood and AIC values to permit direct comparison with other models, we refitted *M*_*cov*=1,*b*=0_ to produce an estimate of apparent abundance without this feature. Specifically, we refitted the model with 20 pseudo-absences in this portion of the study area to better inform abundance-covariate relationships.

The fact that models with and without a PS effect garner approximately equal weight suggests a need to account for PS when producing abundance estimates from this data set. A model averaged estimate calculated using AIC machinery (Burnham & Anderson 2002) is 54854 (SE 9351), which is 7.5% less than the estimate assuming no PS. Notably, the standard error of the model averaged estimate was 79% higher than the model assuming no PS.

## Discussion

In this study, we showed that coarse-grained preferential sampling (Fig. 1) can have a profound impact on the quality of estimates (e.g. animal abundance) when sampling is non-randomized. In simulations, estimators were increasingly positively biased as PS increased. When PS was present, we were able to substantially reduce bias by conducting estimation under a framework where the state variable of interest and the sites chosen for sampling were jointly modeled under a dependent covariance structure. In absence of PS, simulations suggest that this structure results in lower precision than a model without a PS effect; thus the need to account for PS reduces the quality of inference.

Bias attributed to PS may seem counterintuitive, especially given the maxim in survey sampling to allocate more effort to strata for which animal density is high. For instance, in large scale line transect surveys under stratified sampling, the optimal amount of effort that should be allocated to stratum *s* is 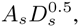, where *A*_*s*_ is the area of *s* and *D*_*s*_ is the anticipated density (Buckland *et al.* 2001; eqn 7.7). Thus, there are theoretical reasons to sample more in high density areas than in low density areas. The obvious solution in this instance is to account for variation in sampling intensity with explanatory covariates or post hoc stratification. However, it is not always clear how to perform post hoc stratification when effort is allocated in a subjective manner.

When applied to bearded seal count data, approximately equal support was given to models with and without a PS effect. The PS effect size was estimated to be positive and to produce considerably lower abundance estimates than models without a PS effect. Differences between apparent abundance estimates decreased when covariates were added to model structure, supporting previous theoretical results (Diggle *et al.* 2010; Pati *et al.* 2011) that covariates serve to decrease the conditional dependence between site selection and the state variable of interest. However, in our data set, adding covariates did not eliminate evidence of PS. Accounting for PS in a model averaging framework led to a moderate decrease in our apparent abundance estimate for bearded seals in this region, and markedly decreased precision. As in our simulations, the need to account for PS thus appeared to have a real cost in terms of variance inflation.

We attempted to fit models to bearded seal data where the degree of PS changed over the landscape, in a similar manner to multivariate spatial models (Royle & Berliner 1999). However, such models led to difficulties with parameter identification in our bearded seal application (see Appendix S2). Evidently, such models may require greater spatial balance or richer data sets. At this time, we suggest limiting initial consideration to models with a single, estimated *b* parameter as a composite adjustment to abundance or occupancy. Although more complex models are clearly identifiable in some situations (Royle & Berliner 1999), further research on the viability of such models as a function of data quality appears warranted. It would also be worthwhile to investigate whether parameter identification varies as a function of the support of the state variable being modeled (e.g. binary vs. count data).

The models we have developed here are specific to spatial models with discrete support, as when data are gathered at a plot level, or aggregated prior to analysis. However, it should be possible to extend our approach to continuous space. One approach would be to model sampling locations as realizations from a spatial point process in a manner similar to Warton & Shepherd (2010). Another possible extension would be to consider models for the sampling process where sampling occurs without replacement for a fixed sample size. For instance, the Bernoulli sampling model makes the implicit assumption that sample size is random. If, instead, a fixed number of locations are sampled, the Bernoulli model is somewhat misspecified. Our simulations suggest some robustness to this misspecification, as the Bernoulli model performed reasonably well when sampling was without replacement for a fixed sample size (Fig. 4). Still, a more precise treatment would need to rely on an extended hypergeometric distribution with variable inclusion probabilities when formulating the sampling model; this extension is nontrivial.

Our conception of PS is related, but not equivalent to “sample selection bias” (e.g. Phillips *et al.* 2009) in presence-only models. In such models, absence of a species at a given site is never directly observed. To draw inference about space use, it is thus necessary to produce a background sample representing the range of locations and habitats that could have been sampled. Sample selection bias then results if the characteristics of sites selected for sampling (e.g. by a volunteer or museum collector) differ systematically from the assumed background sample. In our case, we use PS to refer to the case where absences are available, but where the probability of sampling is dependent on some unknown factor that is also related to abundance or presence of the target species.

## Conclusion

Model-based approaches to estimation of abundance or occurrence have become popular in recent years. We (the authors) have noticed a tendency for analysts to assume that inclusion of spatial covariates or random effects into predictive models will make the underlying sampling design ignorable. We have shown in this paper that this is not the case, although our results do suggest that including predictive covariates can indeed decrease bias from preferential sampling. We have also shown that it is possible to further diagnose and adjust for preferential sampling by jointly modeling dependence between the data collection mechanism and the process of interest (e.g. abundance or occupancy). However, such models can be considerably less precise and have greater instability than models without a preferential sampling parameter. Where possible, we suggest that survey planners incorporate design-based elements (e.g. random or systematic sampling) into their survey designs to reduce the need for model-based triage.

## Acknowledgements

Funding for aerial surveys was provided by the U.S. National Oceanic and Atmospheric Administration and by the U.S. Bureau of Ocean Energy Management. Views expressed are those of the authors and do not necessarily represent findings or policy of any government agency. Use of trade or brand names does not indicate endorsement by the U.S. government.

## Data accessibility

R scripts and data necessary to recreate analyses have been collated into an R package, which is currently available at https://github.com/NMML/pref_sampling. We plan to publish the package to an online archive/repository upon acceptance.

